# Retrograde labeling illuminates distinct topographical organization of D1 and D2 receptor-positive neurons in the prefrontal cortex of mice

**DOI:** 10.1101/2020.05.04.077792

**Authors:** Sara M. Green, Sanya Nathani, Joseph Zimmerman, David Fireman, Nikhil M. Urs

**Author notes:** Correspondence: Nikhil M. Urs, Department of Pharmacology and therapeutics, University of Florida, 1200 Newell Dr, ARB-R5-140, Gainesville FL 32610.

## Abstract

The cortex plays an important role in regulating motivation and cognition, and does so by regulating multiple subcortical brain circuits. Glutamatergic pyramidal neurons in the prefrontal cortex (PFC) are topographically organized in different subregions such as the prelimbic, infralimbic and orbitofrontal, and project to topographically-organized subcortical target regions. Dopamine D1 and D2 receptors are expressed on glutamatergic pyramidal neurons in the PFC. However, it is unclear whether D1 and D2 receptor-expressing pyramidal neurons in the PFC are also topographically organized. We used a retrograde adeno-associated virus (AAVRG)-based approach to illuminate the topographical organization of D1 and D2 receptor-expressing neurons, projecting to distinct striatal and midbrain subregions. Our experiments reveal that AAVRG injection in the nucleus accumbens (NAcc) or dorsal striatum (dSTR) of D1Cre mice labeled distinct neuronal subpopulations in medial orbitofrontal or prelimbic PFC, respectively. However, AAVRG injection in NAcc or dSTR of D2Cre mice labeled medial orbitofrontal, but not medial prelimbic PFC, respectively. Additionally, D2R+ but not D1R+ PFC neurons were labeled upon injection of AAVRG in substantia nigra pars compacta (SNpc). Thus, our data are the first to highlight a unique dopamine receptor-specific topographical pattern in the PFC, which could have profound implications for corticostriatal signaling in the basal ganglia.

**SIGNIFICANCE STATEMENT:** Corticostriatal connections play an important role in regulating goal-directed and habitual behavior, and neuromodulators such as cortical dopamine play an important role in behavioral flexibility. Dopamine receptor expressing D1R+ and D2R+ projection neurons in the cortex mediate the effects of cortical dopamine, but whether these neurons are anatomically organized in a manner that would explain how these neurons mediate these complex effects, is not clear. Our results show a distinct topographical organization of D1R+ and D2R+ PFC pyramidal neurons that project to distinct striatal and midbrain subregions. These results suggest that effects of cortical dopamine are mediated by anatomically localized distinct receptor- and target-defined subcircuits.

## INTRODUCTION

Dopamine regulates normal processes such as motivation, reinforcement-based learning, reward and movement (Palmiter, 2008; Schultz, 2002), and its dysfunction is implicated in many psychiatric and neurological disorders such as schizophrenia, Parkinson’s disease, obsessive compulsive disorder, attention deficit hyperactivity disorder (ADHD), and addiction (Abi-Dargham, 2018; Robinson et al., 2006; Sulzer, 2011; Yager et al., 2015; Zhai et al., 2018).

The striatum is the main input center of the basal ganglia, receiving input from cortical, thalamic, limbic and dopaminergic nuclei (Ikemoto and Bonci, 2014). Glutamatergic signals from cortical areas, and dopamine signals from midbrain dopaminergic nuclei act upon spiny projection neurons (SPNs) in the striatum. The SPNs integrate both glutamate and dopamine signals and coordinate various aspects of learning and behavior (Bamford et al., 2018; Horvitz, 2009; Shiflett and Balleine, 2011).

Afferent cortical glutamatergic inputs into the striatum originate within various medial or lateral subregions of the PFC such as the prelimbic, infralimbic, orbitofrontal, and motor cortex. The PFC plays a critical role in motivation and cognition (Balleine and O’Doherty, 2010; Smith and Graybiel, 2014). Within the medial and lateral PFC, topographically-organized regions such as the dorsally located prelimbic corticostriatal neurons, and the ventral infralimbic or orbitofrontal PFC (OFC) neurons have dissociable effects on motivated behavior and cognitive flexibility (Ahmari et al., 2013; Barker et al., 2017; Hart et al., 2018; Killcross and Coutureau, 2003). (Burguiere et al., 2013; Gremel and Costa, 2013; Rudebeck and Murray, 2011). Although, these studies elegantly outline the role of sub-regions in the PFC or striatum, few studies have explored the neuronal and molecular diversity of PFC pyramidal neurons involved in regulating motivation and cognition.

Dopamine and its receptors in the PFC also regulate motivated behavior and cognitive flexibility (Barker et al., 2013; Goldman-Rakic, 1998; Hitchcott et al., 2007; Natsheh and Shiflett, 2018; Ott and Nieder, 2019). Dopamine activates D1 and D2 class of receptors in the PFC that signal through stimulatory Gαs or inhibitory Gαi proteins, respectively, and through β-arrestins as well, which modulate the activity of both pyramidal neurons and interneurons (Beaulieu et al., 2007; Cousineau et al., 2020; Ferguson and Gao, 2018; Santana et al., 2009; Tomasella et al., 2018; Tseng and O’Donnell, 2004, 2007; Urs et al., 2016). Moreover, pharmacological targeting of D1Rs or D2Rs, or genetic deletion of D2Rs in the PFC can regulate dopamine-dependent behaviors such as locomotion, cognition, and goal-directed behavior (Barker et al., 2013; Del Arco et al., 2007; Hitchcott et al., 2007; Khlghatyan and Beaulieu, 2020; Natsheh and Shiflett, 2018; Tomasella et al., 2018; Urs et al., 2016). Although PFC dopamine receptors play an important role in motivation and cognition, the topographical distribution of D1R+ or D2R+ neurons in specific subregions of the PFC is not known. Here, we use a retrograde AAV (AAVRG)-based approach to identify distinct topographically-organized subpopulations of D1R+ and D2R+ neurons in the PFC based on their target projection areas. Given the role of various subregions of the PFC in motivated behavior and cognition, and the heterogeneity of these regions, the effects of dopamine and its receptors within these subregions will expand our understanding of the molecular and neuronal mechanisms regulating motivated behavior and cognition.

## MATERIALS AND METHODS

### Animals

All mouse studies were performed according to NIH guidelines for animal care and use, and were approved through the University Animal Care and Use Committee. All mice were housed in a 12h light-dark cycle at a maximum of five per cage, provided with food and water *ad libitum* and tested at 8-12 weeks of age. Mice were age matched and mice of both sexes were used, and all experiments were performed in naïve animals. Dopamine D1 receptor Cre (D1Cre, EY262) and Dopamine D2 receptor Cre (D2Cre, ER44) transgenic mice were obtained from MMRRC. Cre+ hemizygous transgenics were used for all experiments.

### Viral Surgeries

D1Cre or D2Cre mice were stereotaxically injected unilaterally with 40-100nl retrograde adeno-associated virus (AAVRG), AAVRG-CAG-Flex-TdTomato (Addgene #238306) and AAVRG-hSyn-DIO-GFP (Addgene #50457). Stereotaxic coordinates are as follows: dSTR (AP +1.0, ML +1.8, DV -3.25), NAcc core (AP +1.0, ML +1.0, DV -4.75), DMS (AP +1.0, ML +1.2, DV -2750), DLS (AP +1.0, ML +2.2, DV -3.25) and SNpc (AP +3.5, ML +1.25, DV -4.1). Mice were allowed to recover for 2 weeks to allow for viral expression of GFP or TdT before imaging and counting of cells.

### Immunostaining, Imaging and Quantification

40 μm thick vibratome cut sections of formalin-fixed mouse brains were processed for imaging. Sections from rostral, rostrocaudal and caudal PFC, and striatal and midbrain sections were imaged using a Nikon AZ100 Zoom microscope, using the same exposure across genotypes for an injection pair. Captured images were used for quantifying number of fluorescent cells for each channel (GFP and TdT) in different subregions of rostral, rostrocaudal and caudal PFC using ImageJ (NIH), and threshold was kept the same between genotypes and injection pairs. At least three sections from rostral, rostrocaudal and caudal PFC were analyzed for each mouse, with an n=5 mice per injection pair. For glutamatergic or GABAergic marker identification, we performed antigen retrieval in citrate buffer at 80 degrees on virally injected PFC sections, and colabeled with antibodies to GFP (Frontier Institute, Japan, Cat# AB_2571575), RFP (Rockland, Cat# 600-401-379), CamKIIa (Enzo Life Sciences, Cat# ADI-KAM-CA002-D), parvalbumin (PV) (Frontier Institute, Japan, Cat # AB_2571613), and GAD 65/67 (Frontier Institute, Japan, Cat# AB_2571698). Imaging for PV and GAD colabeling were done using the Nikon AZ100 zoom microscope. Imaging for CamKIIa labeling was done using a Nikon spinning disk confocal (CSU-X1, Yokogawa) with either 10x or 60x objective on an inverted microscope (Nikon Ti2-E), with a back-thinned sCMOS camera (Prime 95B, Photometrics).

### Statistical analyses

Data were analyzed by a standard two-way ANOVA test for comparison between genotypes, and injection pairs. Individual genotypes, or injection pairs were compared using a *post hoc* Tukey’s test. Data are presented as mean±SEM. p<0.05 is considered as significant.

## RESULTS

For this study we focused primarily on dopamine-related subcortical target regions such as the striatum and midbrain dopamine nuclei i.e. substantia nigra pars compacta (SNpc). The striatum itself can be topographically divided along the dorsoventral or mediolateral axes, into the dorsal and ventral striatum, or dorsomedial (DMS) and dorsolateral (DLS), respectively.

### Dorsoventral Topographical distribution of D1R+ and D2R+ neurons in the PFC

We first injected AAVRGs in topographically distinct target regions along the dorsoventral axis in the dorsal striatum (dSTR) and the nucleus accumbens (NAcc) core within the ventral striatum, respectively, of either D1Cre or D2Cre mice. 100nl of AAVRG-CAG-Flex-TdTomato and AAVRG-hSyn-DIO-GFP were stereotaxically injected in the dSTR or NACC core (see methods for coordinates), respectively. As shown in Figure 1, we observed distinct topographically-organized patterns of GFP (dSTR) and TdT (NAcc core) positive cells in the PFC of D1Cre and D2Cre mice. For D1Cre mice, AAVRG injection in the dSTR (GFP+) and NAcc (TdT+) labeled distinct minimally overlapping subpopulations of dorsally located layer 5, and ventrally located layer 2/3 neuronal cell bodies in the PFC, respectively (Figure 1, ai). The dSTR projecting neurons were primarily localized to dorsally located prelimbic/cingulate (Cg1/PrL) and motor cortex (M1/M2), and mediolaterally located ventral/lateral OFC (VO/LO), whereas the Nacc projecting neurons were primarily localized to the ventrally located medial OFC and infralimbic (MO/IL) regions. Quantifying the number of cell bodies in these regions from rostral, rostrocaudal and caudal PFC sections (see methods) revealed minimal overlap (∼5%) between the PrL and MO/IL subpopulations (Figure 1 c, ** p< 0.01, Two-way ANOVA). “D1-dSTR” - Cg1/PrL: 145 ± 11.3; MO/IL: 22.5 ± 3.9; VO/LO: 121.7 ± 16; M1/M2: 122 ± 11.2 neurons, and “D1-NAcc” - Cg1/PrL: 20.8 ± 9.1; MO/IL: 93.4 ± 10.6; VO/LO: 10.1 ± 2.2; M1/M2: 6.7 ± 6.6 neurons. In contrast, for D2Cre mice, AAVRG injection in the dSTR and NAcc labeled predominantly ventrally located TdT+ layer 5 neuronal cell bodies in MO/IL, but few dorsally located GFP+ neurons in Cg1/PrL (Figure 1, bi). Quantifying the number of cell bodies in these regions revealed minimal overlap of TdT+ neurons (∼15%) between the PrL and MO/IL subpopulations (Figure 1 d, ** p< 0.01, Two-way ANOVA). The NAcc projecting neurons were primarily localized to the ventrally located medial OFC and infralimbic (MO/IL) regions, whereas the few dSTR projecting neurons were primarily localized to M1/M2 and VO/LO regions. “D2-dSTR” - Cg1/PrL: 7 ± 1.8; MO/IL: 3.7 ± 1; VO/LO: 24.4 ± 5.9; M1/M2: 45.7 ± 11.8 neurons, and “D2-NAcc” - Cg1/PrL: 55.7 ± 9.7; MO/IL: 162.9 ± 25.5; VO/LO: 32.8 ± 6.1; M1/M2: 5.6 ± 1.7 neurons. Thus, in the medial PFC of D2Cre mice, MO/IL neurons were predominantly labeled, but not Cg1/PrL neurons.

**Figure 1.**
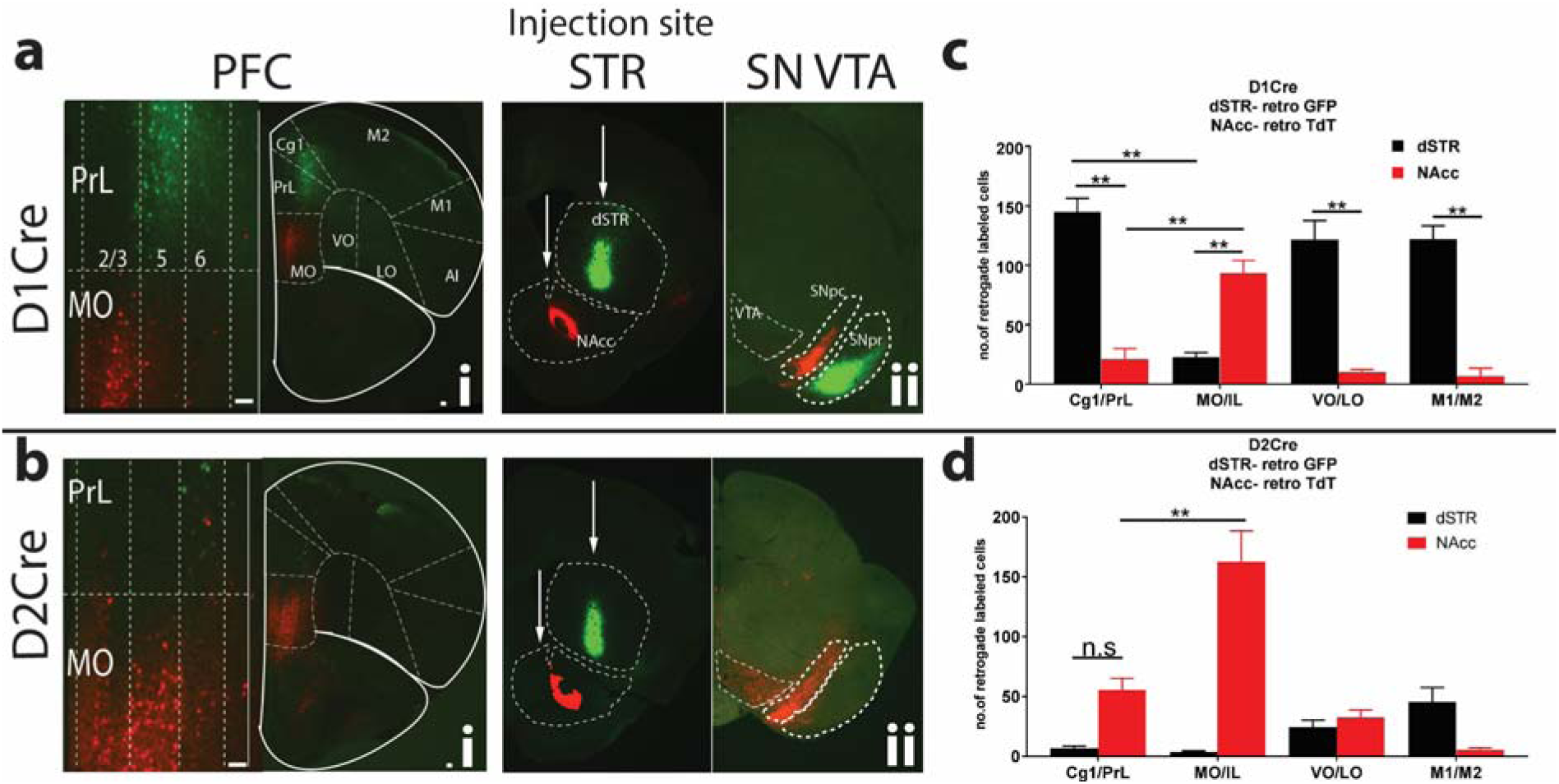
Unique topographically organized subpopulations of D1R+ and D2R+ PFC neurons project to dSTR and NAcc. PFC sections from D1Cre (**ai**) and D2Cre (**bi**) mice showing cell bodies retrograde labeled with both TdTomato and GFP. Left panels are zoomed insets of medial PrL and MO showing layer-specific localization of cell bodies. PrL-prelimbic, Cg1- Cingulate, MO-medial orbitofrontal, VO- ventral orbitofrontal, LO-lateral orbitofrontal, M1- primary motor, M2- secondary motor cortex. **aii, bii.** Injection sites are shown in striatal (STR) sections along with midbrain SNpc/VTA (SN VTA) sections. D1Cre and D2Cre mice were injected with Cre-dependent retrograde AAV, AAVRG-CAG-Flex-TdTomato and AAVRG-hSYN-DIO-GFP in dSTR (100nl) and NAcc (100nl). Representative images (n=5). **c**,**d.** – quantification of # of cells. Data are represented as mean ± SEM. n.s – not significant. ** p<0.01 Two-way ANOVA. Scale bar=100µm.

In both D1Cre and D2Cre mice, distinct projection fibers were observed in midbrain regions i.e substantia nigra pars compacta (SNpc) and reticulata (SNpr). In the D1Cre mice, GFP+ projection fibers (from dSTR) were observed in the SNpr, whereas, TdT+ projection fibers were observed in the SNpc (Figure 1, aii). In contrast, for the D2Cre mice only TdT+ projection fibers were observed in the SNpc, and no labeling in the SNpr (Figure 1, bii).

PFC projection neurons are primarily glutamatergic, but some studies have shown that a small percent of these projection neurons can be GABAergic (Lee et al., 2014; Melzer et al., 2017), and contribute to physiological outcomes. To confirm whether these retrogradely labeled neurons are glutamatergic or GABAergic, we performed colocalization studies with known glutamatergic neuron marker CamKIIa, and known GABAergic neuron markers GAD 65/67 and parvalbumin (PV). As seen in Fig. S1, retrogradely labeled neurons in D1 or D2Cre mice predominantly colocalize with CamKIIa but not with GAD or PV, thus confirming a glutamatergic identity of these corticostriatal projection neurons.

PFC pyramidal neurons also send projections to midbrain regions, specifically to dopamine nuclei (Murugan et al., 2017; Watabe-Uchida et al., 2012). However, whether D1R+ and D2R+ PFC neurons send projections to midbrain dopamine neurons is not known. We injected AAVRGs in dSTR (GFP) as reference, and in SNpc (TdT) of D1Cre or D2Cre mice. For D1Cre mice, similar to Figure 1, injection in the dSTR labeled predominantly distinct Cg1/PrL, VO/LO and M1/M2 localized populations of PFC (Figure 2, ai). However, for SNpc injections we saw minimal labeling of PFC neurons (Figure 2, ai). Quantification of neurons from these regions show a similar predominant labeling of GFP+ neurons in Cg1/PrL as in Figure 1 (Figure 2 b, **p<0.01, Two-way ANOVA), but minimal TdT+ (SNpc) labeling. In contrast, for the D2Cre mice we observed minimal GFP+ labeling similar to Figure 1, but we observed robust TdT+ (SNpc) labeling of PFC neurons (Figure 2, bi). TdT+ SNpc labeling was presumably from dopamine neurons since dense TdT+ axonal projections were seen in the dSTR (Figure 2, bii). Quantification of labeled neurons from these regions revealed however, that there was no topographical pattern for the midbrain projecting D2R+ neurons (Figure 2 d, **p<0.01, Two-way ANOVA). “D2-SNpc” - Cg1/PrL: 22.7 ± 4.3; MO/IL: 17.0 ± 3.4 VO/LO: 39.6 ± 7.1; M1/M2: 29.4 ± 4.5 neurons. Similar patterns were observed for VTA injections for both D1Cre and D2Cre mice as well (Figure S2). Together these data suggest that predominantly D2R+ and not D1R+ PFC neurons project to midbrain dopamine neurons.

**Figure 2.**
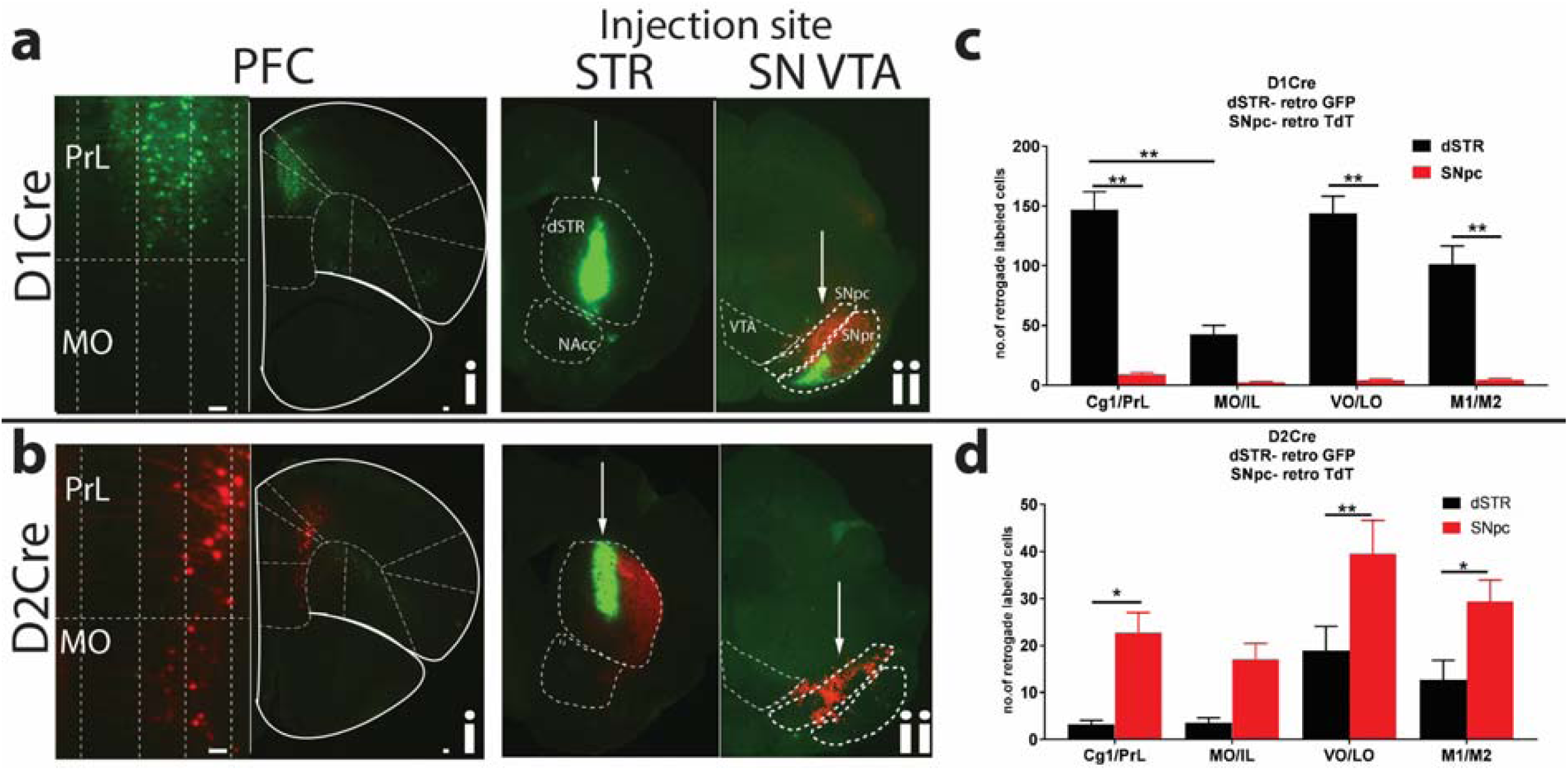
Unique topographically organized subpopulations of D1R+ and D2R+ PFC neurons project to dSTR and SNpc. PFC sections from D1Cre (**ai**) and D2Cre (**bi**) mice showing cell bodies retrograde labeled with either GFP or TdTomato, respectively. Left panels are zoomed insets of medial PrL and MO showing layer-specific localization of cell bodies. PrL- prelimbic, Cg1- Cingulate, MO-medial orbitofrontal, VO- ventral orbitofrontal, LO-lateral orbitofrontal, M1-primary motor, M2- secondary motor cortex. **aii, bii.** Injection sites are shown in striatal (STR) and midbrain SNpc/VTA (SN VTA) sections. D1Cre and D2Cre mice were injected with Cre-dependent retrograde AAV, AAVRG-CAG-Flex-TdTomato and AAVRG-hSYN-DIO-GFP in dSTR (100nl) and SNpc (40nl). Representative images (n=5). **c**,**d.** – quantification of # of cells. Data are represented as mean ± SEM. * p<0.05, ** p<0.01, Two-way ANOVA. Scale bar=100µm.

Figure 3 shows a more detailed comparison of PFC topographical patterns of D1Cre and D2Cre mice for each injection site i.e dSTR, NAcc and SNpc, along the rostro-caudal axis, for the same mice used in Figures 1 and 2. For dSTR (Figure 3a), D1R+ neurons were the predominantly labeled subpopulation across the rostro-caudal axis, in all regions (Cg1/PrL, VO/LO, M1/M2) except in the MO/IL region in rostrocaudal and caudal regions. D2R+ neuron labeling ranged from 1-28% compared to D1R+ neurons for all regions except rostrocaudal and caudal MO/IL. For NAcc (Figure 3b), both D1R+ and D2R+ neurons showed robust labeling across the rostro-caudal axis, with more D2R+ neuron labeling than D1R+ in only the rostral and caudal sections. For SNpc (Figure 3c), D2R+ neurons were predominantly labeled, showing an incremental gradient along the rostro-caudal axis, whereas only 16-20% were D1R+.

**Figure 3.**
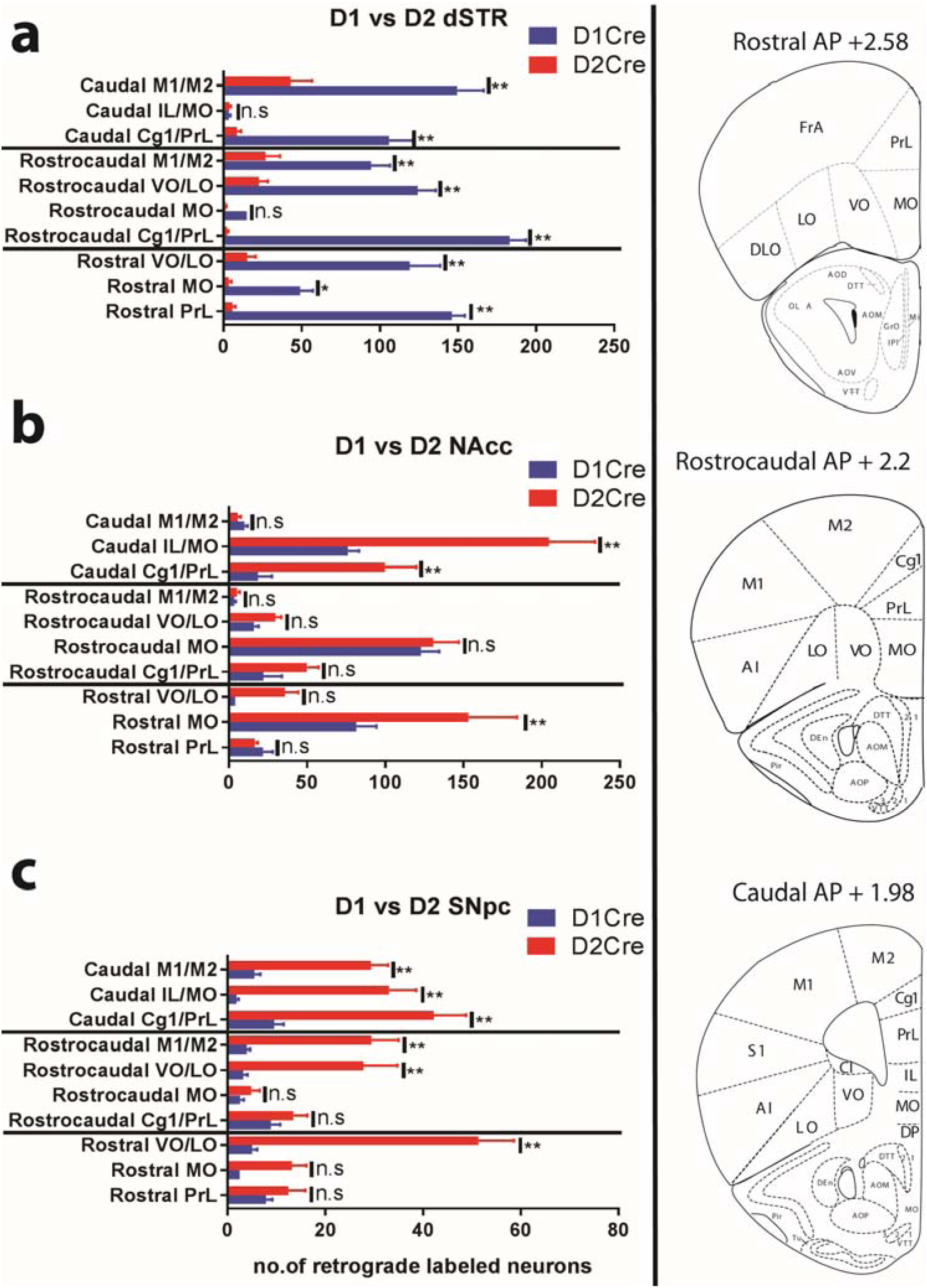
Comparison of topographical distribution of D1R+ and D2R+ PFC neurons along the rostrocaudal axis. **a** – Quantification of GFP+ (dSTR) cells in PFC sections along the rostro-caudal axis of D1Cre and D2Cre mice. Right panel shows representative images and stereotaxic coordinates of rostral, rostrocaudal and caudal PFC sections. **b** - Quantification of TdT+ (NAcc core) cells in PFC sections along the rostro-caudal axis of D1Cre and D2Cre mice. **c** - Quantification of TdT+ (SNpc) cells in PFC sections along the rostro-caudal axis of D1Cre and D2Cre mice. PrL-prelimbic, Cg1- Cingulate, MO-medial orbitofrontal, VO- ventral orbitofrontal, LO-lateral orbitofrontal, AI-agranular insular, M1-primary motor, M2- secondary motor cortex. n=5 for each group. n.s – not significant. Data are represented as mean ± SEM. * p<0.05, ** p<0.01, Two-way ANOVA.

### Mediolateral Topographical distribution of D1R+ and D2R+ neurons in the PFC

In the previous experiments the injection sites were along the dorsoventral axis in the dorsocentral striatum and NAcc core. However, within the dorsal striatum, both medial and lateral subregions have specific roles in motivated behaviors. The dorsomedial striatum (DMS) is involved in the acquisition of goal-directed actions, whereas the dorsolateral striatum (DLS) regulates acquisition of habitual behaviors (Yin et al., 2006; Yin et al., 2005). Previous studies however show that mPFC pyramidal neurons primarily project to the DMS whereas more posterior lateral sensorimotor cortex neurons project to the DLS (Kupferschmidt et al., 2017; Shiflett and Balleine, 2011).

We next asked the question whether PFC D1R+ and D2R+ neurons have specific projection pattern to the DMS or DLS. Similar to Figures 1 and 2, we injected 50nl of Cre-dependent AAVRG GFP and AAVRG TdT in the DMS and DLS, respectively, of D1Cre and D2Cre mice. As seen in Figure 4, we observe distinct topographical patterns for D1R+ and D2R+ neurons projecting to DMS and DLS. For DMS injection in D1Cre mice, we observed robust GFP+ labeling predominantly in the PrL (67 ± 12.5 neurons), whereas for DLS injection we observed robust TdT+ labeling predominantly in M1/M2 (118.3 ± 12 neurons) and AI (146 ± 24.1 neurons) (Figure 4, a and c). Overall, very few D2R+ neurons project to either DMS or DLS (Figure 4, b and d). Comparison of patterns of D1Cre and D2Cre mice for DMS show that only Cg1/PrL has significantly greater labeling of D1R+ neurons (Figure 4e, **p<0.01, Two-way ANOVA). For DLS however, Cg1/PrL, AI and M1/M2 show significantly greater labeling for D1Cre compared to D2Cre mice (Figure 4f, **p<0.01, Two-way ANOVA).

**Figure 4.**
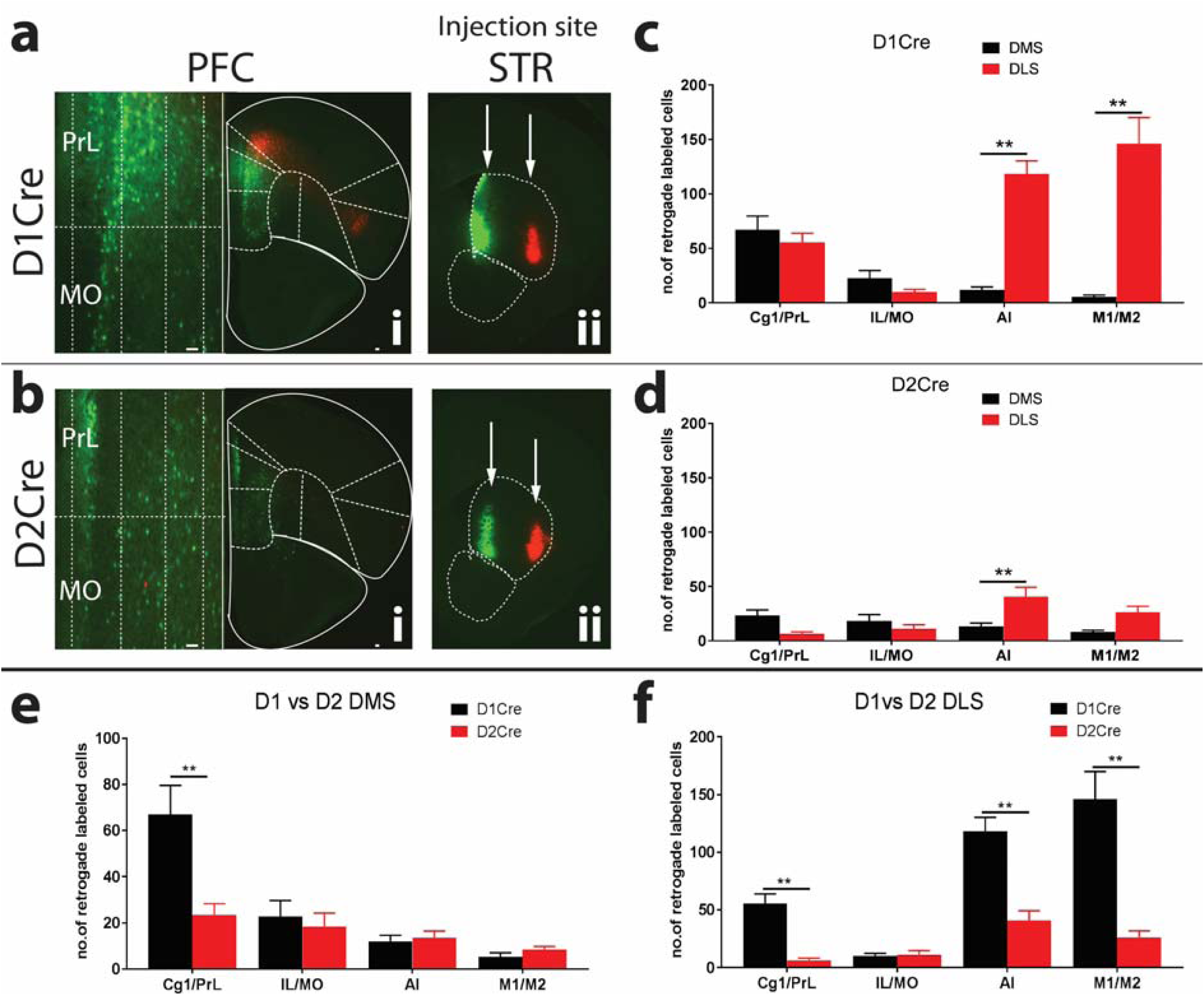
Topographical organization D1R+ and D2R+ neurons projecting to DMS and DLS. PFC sections from D1Cre (**ai**) and D2Cre (**bi**) mice showing cell bodies retrograde labeled with either GFP or TdTomato, respectively. Left panels are zoomed insets of medial PrL and MO showing layer-specific localization of cell bodies. PrL-prelimbic, Cg1- Cingulate, MO-medial orbitofrontal, AI – agranular insular, M1-primary motor, M2- secondary motor cortex. **aii, bii.** Injection sites are shown in DMS and DLS subregions in striatal sections. D1Cre and D2Cre mice were injected with Cre-dependent retrograde AAV, AAVRG-CAG-Flex-TdTomato and AAVRG-hSYN-DIO-GFP in DMS (50nl) and DLS (50nl). Representative images (n=5). **c**,**d.** – quantification of # of cells from D1Cre and D2Cre mice. **e, f** – Comparison of # of labeled cells in PFC of D1 and D2Cre mice projecting to DMS and DLS subregions. Data are represented as mean ± SEM. * p<0.05, ** p<0.01, Two-way ANOVA. Scale bar=100µm.

## DISCUSSION

D1R+ and D2R+ neurons are found throughout all PFC subregions (Anastasiades et al., 2019; Khlghatyan and Beaulieu, 2020; Santana and Artigas, 2017; Yu et al., 2019), but this widespread expression pattern does not adequately explain how these neurons mediate distinct physiological and behavioral outcomes. In this study, we show a previously unappreciated distinct topographical organization of D1R+ and D2R+ neurons in the PFC of mice. As summarized in Figure 5, we observe distinct topographical organization patterns of D1R+ and D2R+ neurons along the dorsoventral and mediolateral axes, based on their projection target. Along the dorsoventral axis, PFC D1R+ neurons are topographically organized such that D1R+ neurons in prelimbic regions primarily project to the dSTR, whereas D1R+ neurons in medial OFC and infralimbic regions primarily project to the NAcc core. In contrast, PFC D2R+ neurons have a distinct pattern of organization, such that very few prelimbic D2R+ neurons project to dSTR, but medial OFC and infralimbic D2R+ neurons project to NAcc core, similar to D1R+ neurons. Along the mediolateral axis, D1R+ neurons in the medial prelimbic region primarily project to dorsomedial striatum, whereas D1R+ neurons in M1/M2 motor and agranular insular cortex primarily project to dorsolateral striatum. In contrast very few D2R+ neurons project to either DMS or DLS. However, medial PFC D2R+ but not D1R+ neurons project to midbrain dopamine nuclei. Thus, our data provide, for the first time a detailed insight into the anatomical organization of D1R+ and D2R+ neurons in the PFC.

**Figure 5.**
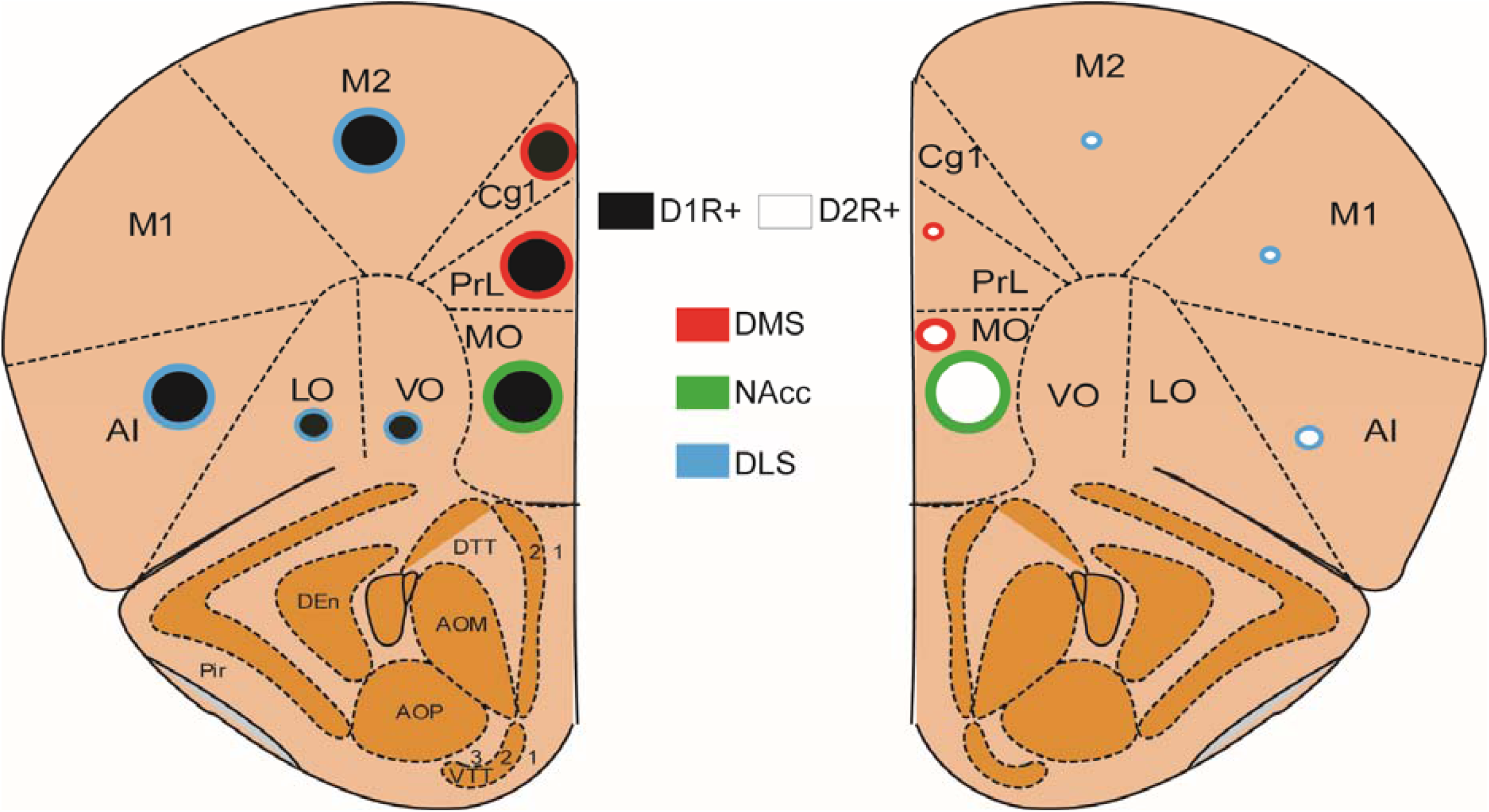
Summary of topographical organization of D1R+ and D2R+ pyramidal neurons in mouse PFC. D1R+ neurons represented by black circles and D2R+ neurons are represented by white circles. Circle borders represent corticostriatal projection targets. Size of the circle represent abundance of projections to target regions.

In this study we use AAVRGs to identify afferent PFC inputs into various striatal and midbrain regions. One potential caveat with using AAVRGs is that we cannot control for variability of infection at the injection site, even if we inject the same volume of AAV. However, one advantage of using AAVRGs is that target neurons can be specifically labeled in a Cre-dependent manner, and therefore label cell bodies of specific populations of neurons using transgenic Cre mice. Our data are consistent with previous findings that D1R+ neurons are primarily corticostriatal, whereas D2R+ neurons are corticostriatal and also project to more caudal regions such as the thalamus (Gee et al., 2012). Other groups have also shown that PFC D1R+ and D2R+ neurons also project to other limbic areas such as the basolateral amygdala (BLA) (Jenni et al., 2017). PFC neurons projecting to BLA, NAcc or VTA are not only distinct subpopulations but also have distinct laminar distribution (Murugan et al., 2017). In this study we see distinct laminar distribution of both D1R+ and D2R+ PFC neurons. “D1-dSTR” prelimbic neurons are predominantly localized to layer 5, whereas “D1-NAcc” MO/IL neurons are predominantly localized to layer 2/3. Interestingly, “D2-NAcc” MO/IL neurons are predominantly localized to layer 2/3 and 5.

Our data also suggests that distinct predominantly D1R+ neuron subpopulations along the mediolateral axis (Cg1/PrL vs M1/M2/AI), project to the DMS and DLS, respectively. However, D2R+ PFC neurons do not project to either DMS or DLS. Thus, D1R+ PFC neurons might be directly involved in regulation of switching between goal-directed versus habitual actions (Yin et al., 2006; Yin et al., 2005). Interestingly, a similar topographical pattern is maintained within the SNpc, where medial and lateral dopamine neurons project to DMS and DLS, respectively (Lerner et al., 2015).

The dSTR-NAcc injections (Figure 1) reveal distinct projection fiber patterns in the SNpr and SNpc. For injections in dSTR, we observed projection fibers in the SNpr only in D1Cre and not D2Cre mice, consistent with AAVRG-GFP labeling striatal D1R+ direct pathway SPNs. However, for NAcc core injections, we observe projection fibers in the medial SNpc and not the SNpr in both D1Cre and D2Cre mice. Our findings are consistent with previous observations that D1R+ and D2R+ NAcc core SPNs do not follow the traditional direct/indirect dichotomy like dSTR SPNs, and instead send projections to ventral pallidum (VP) or midbrain (Kupchik et al., 2015; Pardo-Garcia et al., 2019; Sesia et al., 2014). Moreover, these studies suggest that both D1R+ and D2R+ NAcc core SPNs project to VP, but only D1R+ NAcc core SPNs project to the midbrain. Although we observe projection fibers in the SNpc of D2R+ mice, these are likely direct projections from the MO/IL PFC (Figure 2b), and not from NAcc core SPNs. Thus, D1R+ pyramidal neurons in the MO/IL projecting to NAcc core, can not only release glutamate and regulate excitability of GABAergic SPNs, but also modulate dopamine release in the DMS by indirectly acting on medial SNpc dopamine neurons. In contrast, D2R+ pyramidal neuron in the MO/IL project directly to both NAcc core and SNpc. One possible caveat of this interpretation is that by using AAVRGs we are unable to establish whether these fibers in the SNpc are afferents on GABAergic or dopaminergic neurons. A more sophisticated approach with rabies virus retrograde labeling, with Cre-dependent labeling of target neurons is required to confirm our interpretation.

The dorsal striatum is topographically divided into the DMS and DLS, which have been implicated in action-outcome learning and stimulus-response learning, respectively, whereas, the NAcc, has been implicated in reward perception (Shiflett and Balleine, 2011). D1R+ and D2R+ neurons in topographically organized regions in the PFC can thus have various effects on physiology and behavior depending on their striatal or midbrain projection target, and modulation by cortical dopamine.

Illuminating this unique pattern of organization of D1R+ and D2R+ neurons in the PFC will help us better understand the regional and global effects of cortical dopamine and dopamine receptors in the regulation of motivation and cognition.

---

**Figure S1.**
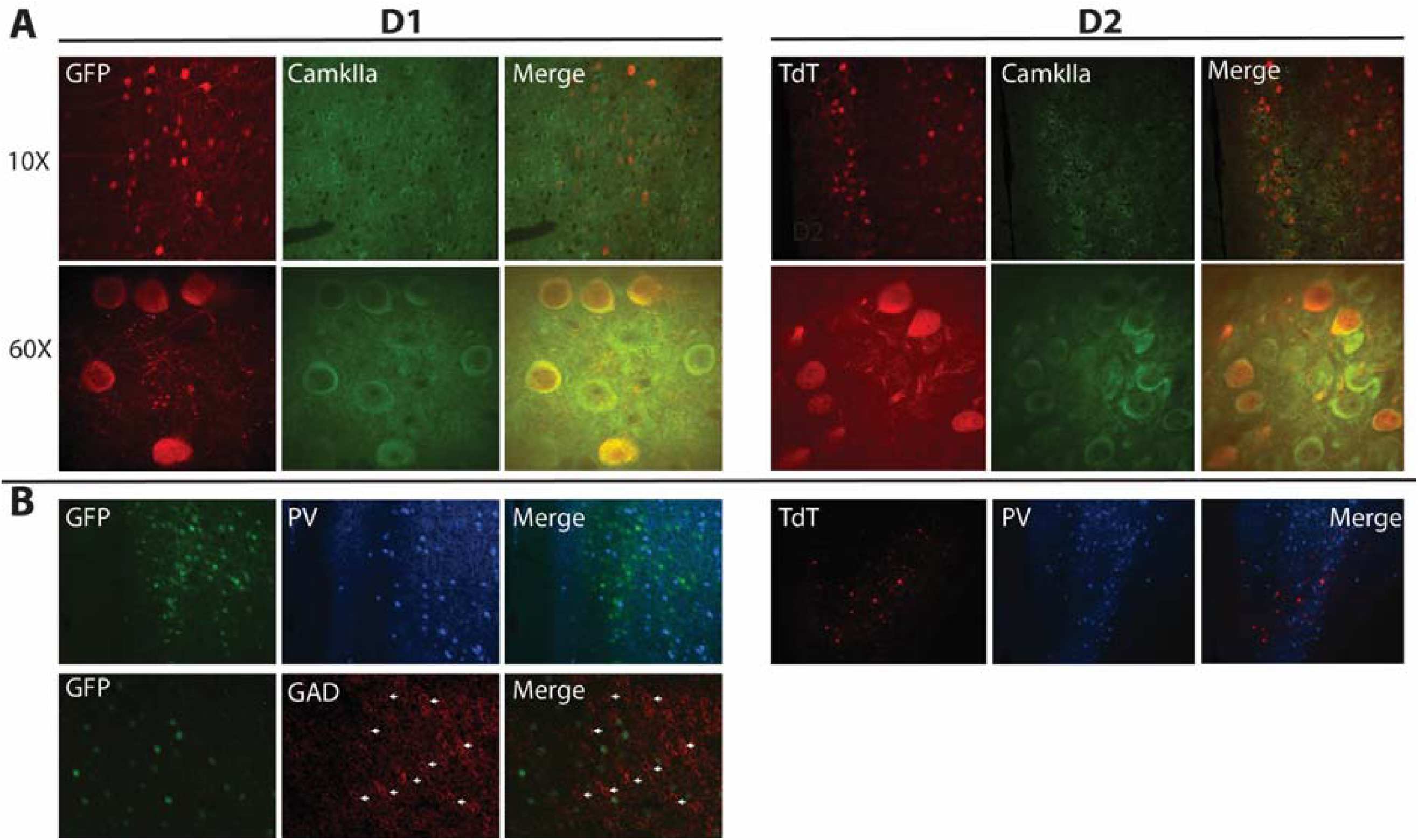
Colabeling of D1R+ and D2R+ PFC neuron subpopulations with glutamatergic and GABAergic markers. PFC sections from D1 and D2Cre mice (n=2) previously retrogradely labeled with GFP or TdT were colabeled for CamKIIa (**A**) or PV and GAD 65/67 (**B)**.

**Figure S2.**
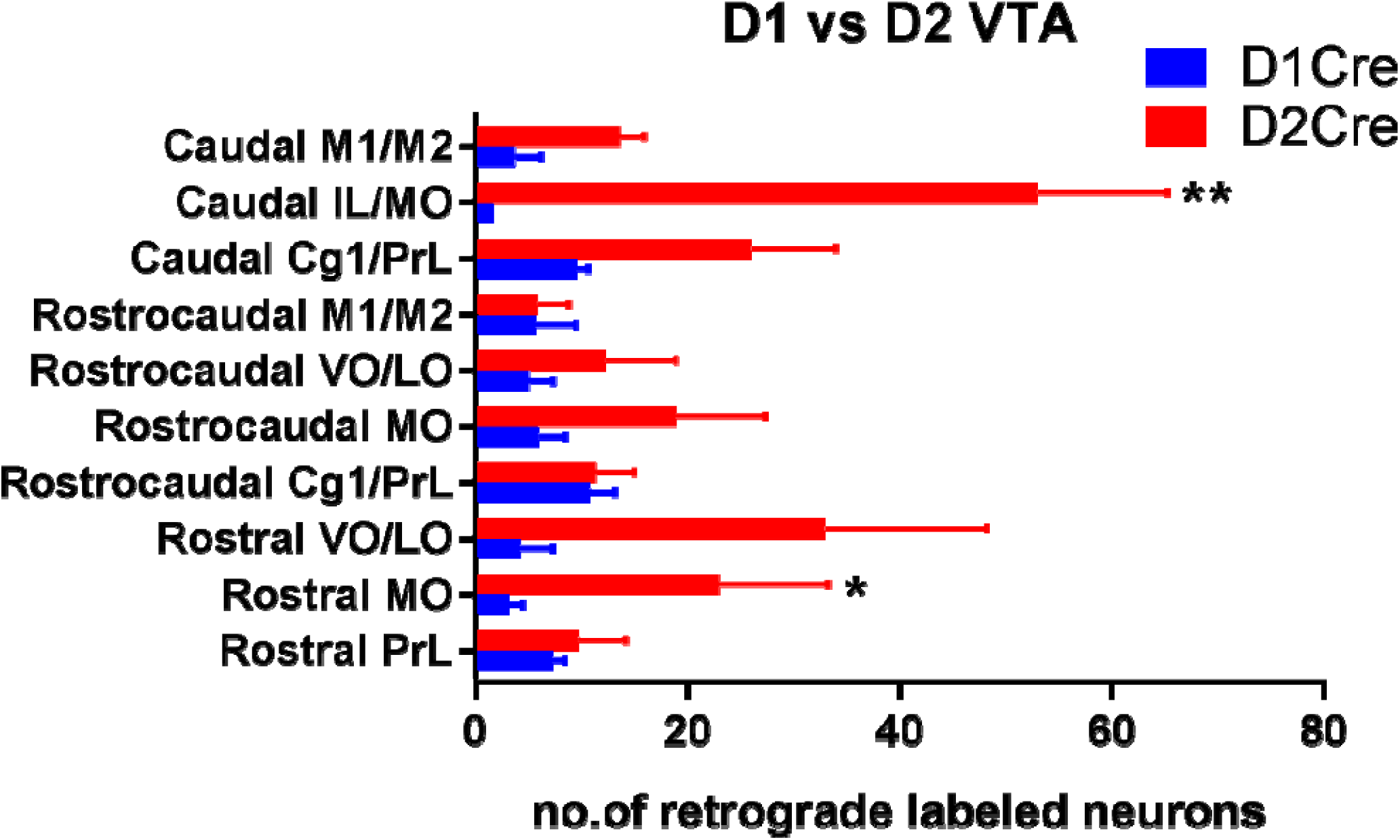
Topographical distribution of VTA projecting D1R+ and D2R+ PFC neuron subpopulations. Quantification of GFP+ (VTA projecting) cells in PFC sections along the rostro-caudal axis of D1Cre and D2Cre mice. PrL-prelimbic, Cg1- Cingulate, MO-medial orbitofrontal, VO- ventral orbitofrontal, LO-lateral orbitofrontal, M1-primary motor, M2- secondary motor cortex. n=3 for each group. Data are represented as mean ± SEM. * p<0.05, ** p<0.01, Two-way ANOVA.

