## Supplementary figures and images for "Retrograde labeling illuminates distinct topographical organization of D1 and D2 receptor-positive neurons in the prefrontal cortex of mice"

### Supplemental Figure 1

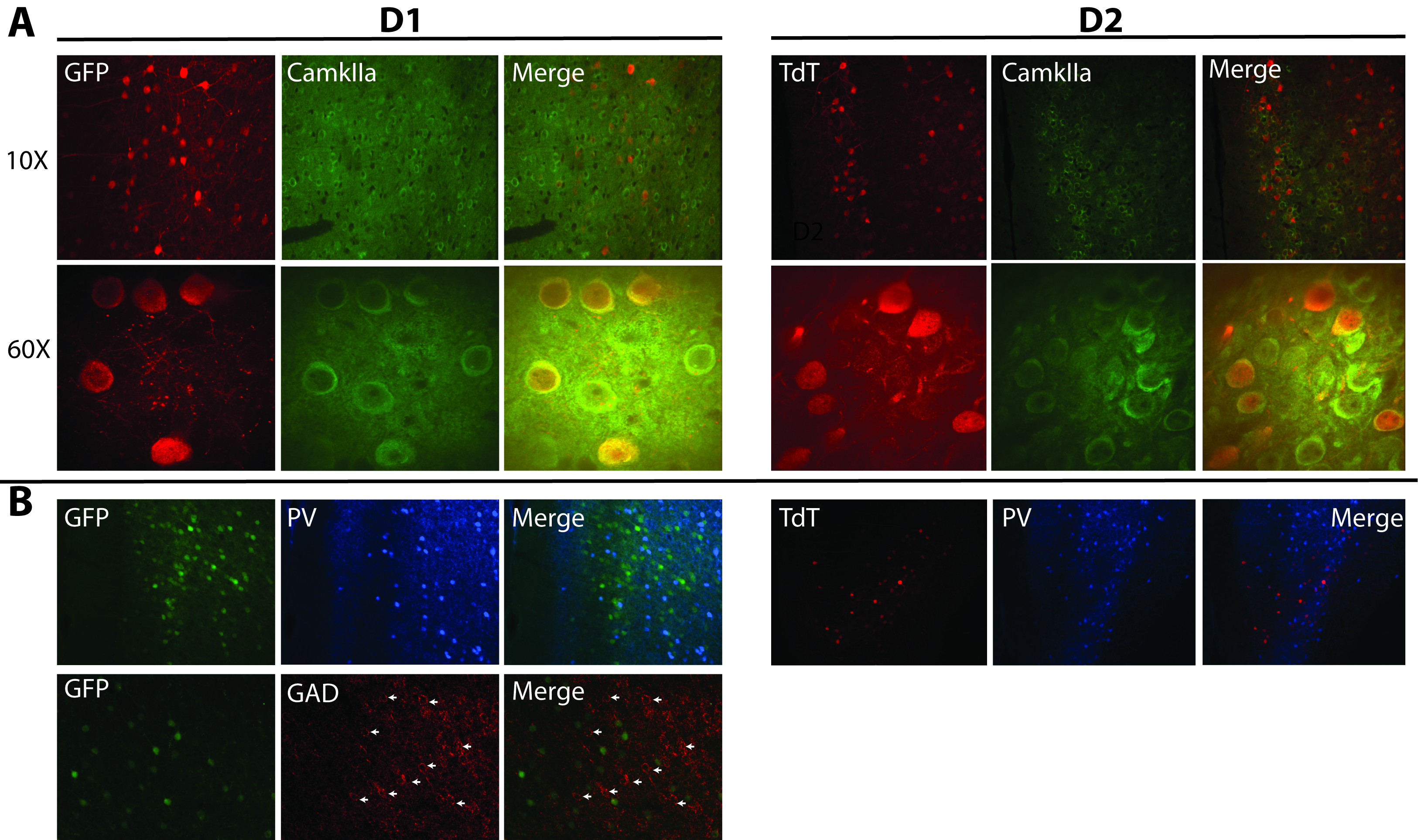

### Supplemental Figure 2

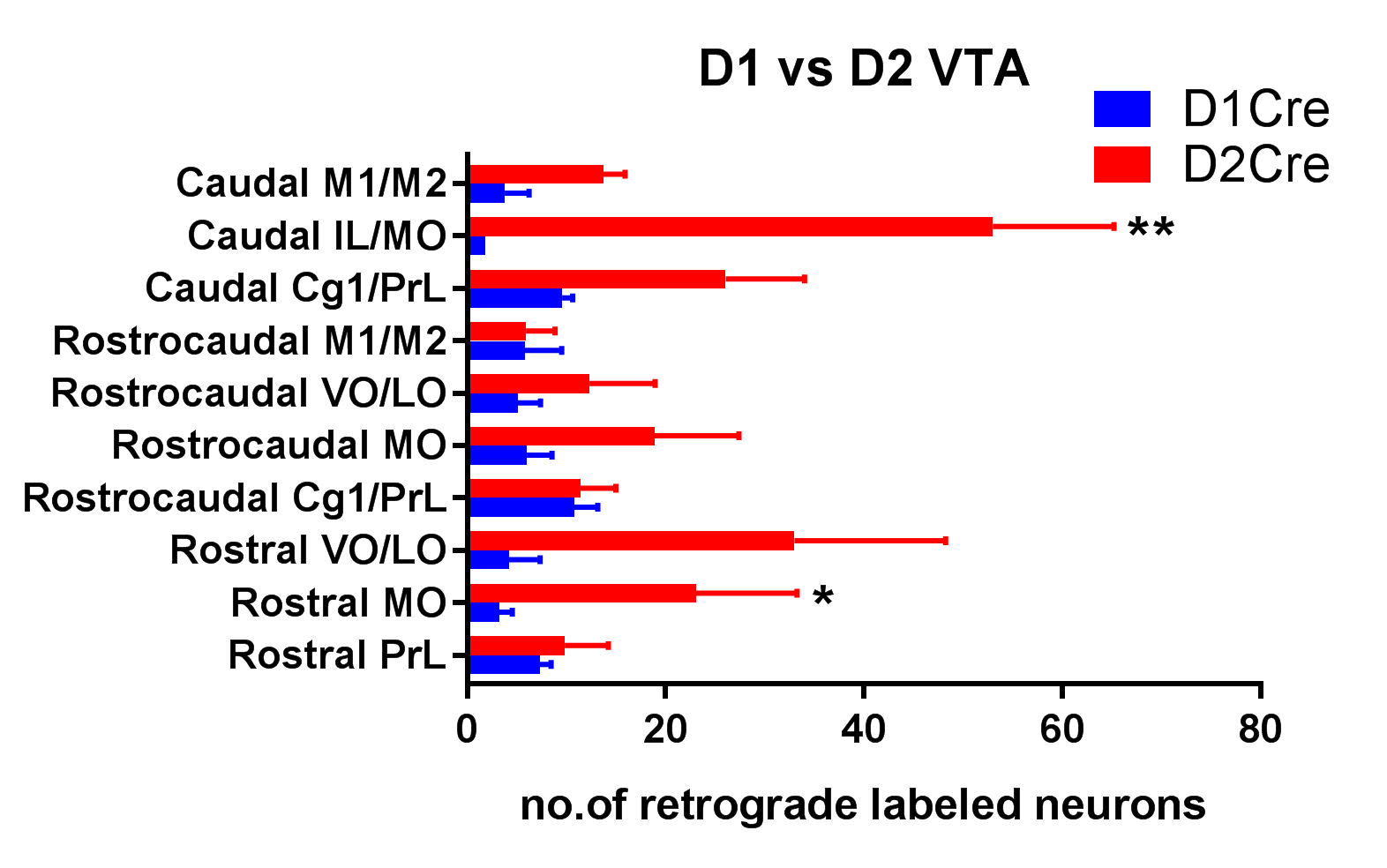
